# Representation of Task Structure in Human Hippocampus and Orbitofrontal Cortex

**DOI:** 10.1101/794305

**Authors:** Eda Mizrak, Nichole R. Bouffard, Laura A. Libby, Erie Boorman, Charan Ranganath

## Abstract

The hippocampus is thought to support episodic memory, or memory for specific events, but recent work also suggests that it may be involved in extracting structure from the world to guide future decisions and predictions. Recent evidence in rodents suggests that the hippocampus supports decision-making in cooperation with orbitofrontal cortex (OFC), possibly based on representation of task structure. Here, we used functional magnetic resonance imaging (fMRI) to test how the human hippocampus and OFC represents decision-relevant information extracted from previous experiences. Participants performed a task in which they learned values of different foods in grocery store contexts. The task was structured such that we could examine the degree to which neural representations could reflect generalized information about different task structures. Specifically, we manipulated whether a food’s desirability varied with store context or not. Some foods were desirable in some store contexts and not in others; some foods were always desirable or undesirable. Participants needed to extract these two task sub-structures (i.e., context-determined vs. context-invariant) from the task structure. We examined hippocampal and OFC activity patterns during a decision-making task after participants were trained with the task structure. Our results showed that both hippocampus and OFC carried task structure information that was relevant to the decision outcomes. Hippocampal and lateral OFC representations differentiated between context-determined (deterministic) and context-invariant (probabilistic) task structures. The degree of this differentiation, an index of task structure representation, was highly correlated between hippocampus and lateral OFC. These results add to a mounting evidence suggesting that the hippocampus and OFC support decision-making by representing task relevant information to the decision outcomes after the task structure is learned.

## INTRODUCTION

Our decisions often rely on our past experiences. For instance, in your local grocery store, you might skip the produce section because previously the produce did not live up to your expectations. Humans can efficiently extract regularities across experiences to make adaptive decisions (i.e., not buying the produce), and several possible mechanisms have been proposed to explain this process.

One simple possibility is that people draw from specific experiences (i.e., the banana that I bought last week at the supermarket was not good). Neuroscience research has reliably shown that the hippocampus is critical for episodic memory formation and retrieval (Moscovitch et al., 2016; Ranganath, 2010). It has been suggested that hippocampus bridges past experiences with future decisions (Biderman et al., 2020). According to some models, the hippocampus binds and represents item and context information from events and episodes such as banana and local grocery store. The binding of items in context, or creating strong associations between events and their spatial or non-spatial contexts, has been widely studied for animal and human hippocampus (Davachi, 2006; Manns & Eichenbaum, 2009; McKenzie et al., 2014; Ranganath, 2010). A related view is that the hippocampus might form chains of overlapping associations (Horner et al., 2015; Schlichting & Preston, 2015) or statistical regularities (Schapiro et al., 2016, 2017) that integrate across multiple events.

Another non-exclusive possibility is that we might form more general representations that capture the abstracted structure of an environment or task, and so that we can make future decisions in a flexible manner. For instance, when shopping, you might have a structured representation or “cognitive map” (Tolman, 1948) of your neighborhood about which store provides the best produce and which stores to skip. Later, we use this cognitive map to guide grocery shopping decisions. One theory proposes that the hippocampus might encode the cognitive maps that might be generally deployed during decision-making (O’Keefe & Nadel, 1978). Although the concept of cognitive maps has been historically applied to maps of physical space, recent work has emphasized a more general role for the hippocampus, in capturing the structure of a task, or a “task space” that transcends spatial contexts (Behrens et al., 2018; Bellmund et al., 2018; Schiller et al., 2015). A task space consists of information about the interrelationships between all the factors that are relevant to a decision for attaining the desired goal. For instance, the hippocampus could form structured representations of task-relevant information that has been learned/generalized across past experiences, task structure representation.

In addition to the hippocampus, orbitofrontal cortex (OFC) is another region that supports decision making (i.e., not buying a banana at local grocery store) by drawing on outcomes of previous experiences (see reviews by Rushworth et al., 2011; Wallis, 2007). According to this view, OFC represents the predicted and experienced value of behavioral outcomes and is responsible for signaling this value to other regions to help guide current or future decisions (Murray & Rudebeck, 2018; Padoa-Schioppa & Conen, 2017; Rushworth et al., 2011; Wallis, 2019). For example, the next time we are the grocery store, the OFC can provide value prediction for bananas which can be used to predict the outcome and to guide a shopping decision. A recent theory, however, suggests that OFC might have a more general function beyond representing value, and additionally the OFC might represent the structure of the task or environment (Wilson et al., 2014). For instance, studies provided evidence for OFC representing the current task state in a cognitive map of the task spaces (Park et al., 2020; Schuck et al., 2016; Wikenheiser & Schoenbaum, 2016; Zhou et al., 2021). In this view, task structure representation is a more general role for OFC that also includes value representation that can be derived from the task structure.

Recent studies also unpacked the relationship between hippocampus and OFC in representing task structure. For instance, it has been suggested that the hippocampus might encode concrete and detailed information about the environment. This information can be spatiotemporal context or incidental associations between events which is then received by the OFC as an input and tailored for the cognitive needs, extracting the most relevant information for decision-making. Additionally, Wikenheiser et al. (2017) showed that when hippocampal inputs to OFC were suppressed, OFC encoding of the latent context – a critical part of the reversal task structure - was impaired. This finding suggested that OFC representation of the task structure is shaped by the inputs received from the hippocampus. In sum, it has been proposed that both the Hippocampus and OFC encode the cognitive map of a task space; the task structure (reviewed in Wikenheiser & Schoenbaum, 2016).

The current study examined the role of hippocampus and OFC in representing task related information. We used representational similarity analysis (RSA) of functional magnetic resonance imaging (fMRI) data to investigate what type of information the hippocampus and OFC represented in a memory-guided decision-making task. Participants learned about desirability of foods in different stores where a latent task structure was available. Specifically, participants learned about customer preferences for eight food items in four different grocery stores. After learning the customer preferences in each store, participants were scanned while they decided whether a food was desired or not by customers based on previous learning for each food in each store context.

The task was designed such that, desirability of some of the foods were deterministic, either “liked” or “disliked” depending on the specific store context (See Fig. 1). These foods are referred to as “context-determined” (CD) foods, because information about the context was needed in order to make accurate decisions about the customer’s preference (i.e., probability of being “liked” = 1.0 or 0 depending on the store). The desirability of other foods was probabilistic, and this probability did not change across stores (i.e., probability of being “liked” = 0.75 or 0.25 in every store). These foods are referred to as context-invariant (CI) because the context was irrelevant to decisions about the preferences. This difference between the foods introduced two distinct sub-structures of the task. In the case of our study, these two sub-structures determined whether food outcomes depended on the store context or not. Participants were not explicitly informed about these two sub-structures; they needed to learn this abstract information from the distribution of like/dislike probabilities across stores and foods. In other words, the task structure was hidden or latent because they needed to be inferred from the food outcome distributions. If participants were able to learn these two sub-structures, they could generalize over them and lump these foods together; representing them more similarly to each other. In addition to the latent task structure, some foods had similar outcomes across stores, or some stores provided similar outcomes across foods (see Figure 1B). The study design enabled us to observe how participants learn a simple latent task structure and how task related information is represented and retrieved in the hippocampus and OFC when participants are making decisions.

**Figure 1.**
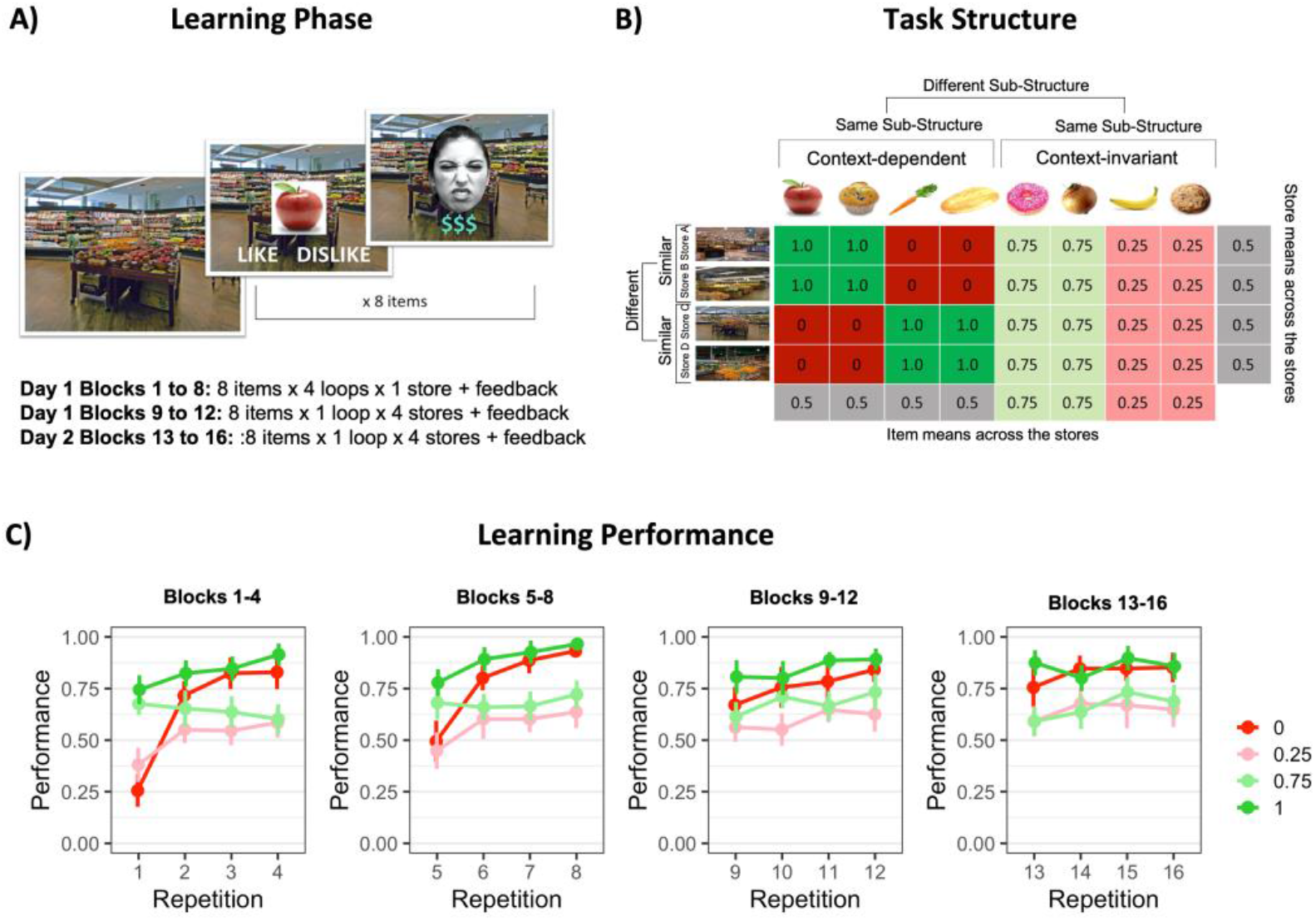
**A) Learning Phase.** Illustration of a trial from the learning phase. On Day 1 participants learned the customer preferences for each food in each store. For the first 8 blocks, subjects are in one store for the entire block and loop through all 8 foods four times. In blocks 9-12, subjects loop through all four stores within a single block, looping through all 8 foods once in each of the four stores. For each trial, one food item is shown on the store context and the participant is asked to predict if customers like this food item in this store context or not. After making their prediction, the participant was given feedback and were told whether they were correct or not along with a picture of a customer’s reaction to that food. At the end, participants would have seen each food item X times in each store. **B) Task Structure.** During the learning phase, participants made predictions about food preferences, and then they received feedback about the actual outcome; customer preference for that food in that store. The matrix illustrates the distribution of outcomes for eight different foods (columns) in each of the four stores (rows). Each cell corresponds to the probability of a like outcome for the given food within the given store; 1.0 means 100 % liked, 0 means 0 % liked or 100 % disliked. Context-Determined foods are defined as those with outcomes that are determined by the store context (left side of the matrix) and Context-Invariant foods are those with probabilities that are the same across stores (right side of the matrix). The task was designed so that there two task sub-structures; context-determined and context-invariant. Additionally, there were pairs of similar contexts for which the same foods were liked and disliked in both stores. **C) Learning Performance.** Performance across all 16 blocks of learning. Foods were separated into four categories based on their like outcome probabilities: 1, 0, .75, .25. A repetition consisted of four loops of learning of each food in each store. Therefore, first repetition was the first time a food category was learned at each store. Learning performance was calculated for each food category by averaging across trials of the foods at each store. Performance corresponded to the proportion of responses of the most likely outcome for a food in a store; like responses for foods with like probabilities of 0.75 and 1, dislike responses foods with like probabilities of 0.25 and 0. Performance increased with repetitions and food preferences for each food category were learned with above 60 % performance at the end of the learning blocks. Error bars denote within-subjects 95 % confidence intervals.

One possibility is that the hippocampus represents each individual event with a unique neural pattern (item + context). Another possibility is that shared item (i.e., food) and context (i.e., store) information is represented in a structured way that can be observed as a neural pattern generalization across these shared dimensions. The OFC might behave similarly to the hippocampus and represent foods based on their outcomes in a structured way and could generalize across foods with similar outcomes (Zhou et al., 2019). Accordingly, there were different types of information that could be extracted from the overarching task structure; the sub-structures (CD vs. CI), item, and context-specific information. We investigated the representational structure of human HC and OFC, and their relationship, while people made context-dependent and context-invariant decisions.

## RESULTS

### Learning phase: Participants learned desirability of foods in stores

Participants first learned about the implicit relationships between the foods and store contexts before they used this knowledge to make decisions about the foods (See Fig. 1A). Participants were not explicitly instructed about the task structure (Fig. 1B) that determined the probabilities of customer preferences for each food. They learned desirability of foods by trial and error based on the feedback they received. Participants completed 12 blocks of training with feedback on decision outcomes on Day 1. After this training phase, we assessed participants’ performance. Only participants who performed above a criterion performance of 60 % optimal predictions for both CD and CI foods were admitted to the second part of the study in the scanner.

We hypothesized that learning performance for context-determined vs. context invariant foods would be different for two reasons; 1) CD foods required learning of the importance of context for the outcome and generalization across contexts would be harmful for these foods whereas for CI foods, context was not a decision-relevant variable, and 2) CD foods were deterministic and CI foods were probabilistic. These differences across the two structures could result in two types of decision-making; a) deterministic, context-dependent, b) stochastic, context-invariant. If the participants differentiated across these two task structures, they could perform differently for CD and CI foods.

To assess learning differences between the two sub-structures, we scored each decision during the learning phase as a function of whether the participant made the *optimal* decision for that trial type. On CI trials, the optimal decision was defined as the most likely outcome for that food item. If a food was liked on 75% of trials, “like” predictions were scored as optimal, whereas if it was liked on only 25% of the trials, then “dislike” predictions were deemed as optimal. In other words, even without any knowledge of the task structure, participants could make accurate predictions solely from learning the relative frequencies of “like” and “dislike” predictions for that food item. For example, in the example depicted in Figure 1B, it would be optimal to choose “Dislike” for the banana and “Like” for the donut, regardless of the current store context.

On CD trials, the optimal prediction was based entirely on the store context. For example, with the information depicted in Figure 1B, it would be optimal to decide that the apple is liked in Stores A and B and disliked in Stores C and D. Across all trials, the “like” probabilities for all CD foods summed to 50%. Accordingly, outcomes on CD trials could only be accurately predicted if the participant learned to adopt context-dependent decisions.

We calculated proportion of optimal responses for foods with different outcome probabilities (CD foods: 0 and 1, CI foods: 0.25, 0.75) and for each repetition averaging across stores. Figure 1C shows learning performance over the course of training. We only analyzed the data from Blocks 13 to 16 on Day 2 after learning had occurred and performance stabilized with a repeated measures ANOVA including sub-structure (CD vs. CI), repetition (13 to 16), and valence (like vs. dislike). As can be seen in Fig. 1C, the proportion of optimal predictions was significantly higher on CD trials than on CI trials (main effect of sub-structure: *F*(1,21) = 31.18, p < 0.001).

### Decision Phase: Participants used their knowledge of the task structure when making decisions about foods

After completing the learning phase, participants were asked to decide whether a food was liked or not in a store context and were scanned during the decision phase. As in the Training Phase, on each trial commenced with presentation with a food image overlaid on a store context, and the participant was instructed to predict the customer preference (either like or dislike) for the presented food in the given store (Figure 2A). Critically, during this phase, <u>no</u> feedback was given, so participants needed to rely on what was learned during the Training Phase to optimally predict customer preferences for each food.

**Figure 2.**
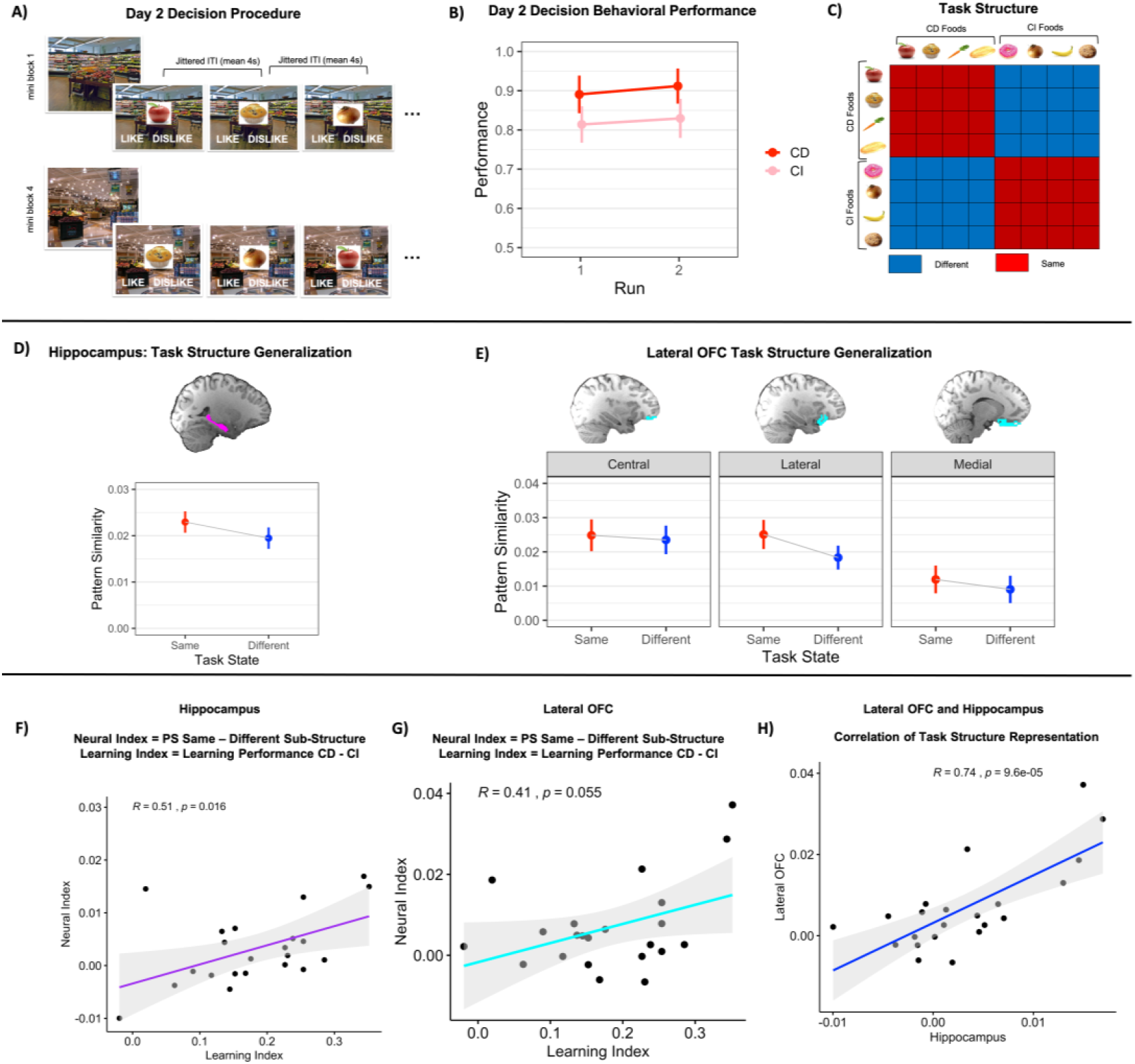
**A) Decision Phase.** Trial procedure was like learning blocks 9 to 16 with the only difference being that no feedback was given after decisions. Same eight foods were shown in each mini-block in different contexts but in a randomized order. Each run consisted of 8 mini-blocks. **B) Decision Performance.** Foods were categorized as context-dependent (CD) and context-invariant (CI) and the proportion of optimal decisions was calculated for CD and CI foods separately and averaged across stores for each run. Decision performance showed that participants learned the task structure well enough to make optimal decisions for both CD and CI foods. CD foods were slightly better than CI foods (p = 0.049). **C) Task structure.** We calculated pattern similarity (PS) for the two same sub-structure trial pairs (CD-CD or CI-CI) and different sub-structure trial pairs (CD-CI). **D) Task structure Representation in Hippocampus.** Behavioral performance indicated participants learned the difference between CD and CI task structures. Pattern similarity for same task structure trial pairs was higher than different task structure (*p* = 0.038). **D) Task structure Representation in OFC.** Lateral OFC pattern similarity was higher for same task structure compared to different task structure. **E) Learning and Task structure in. Hippocampus.** We calculated a neural index for task structure representation in hippocampus, PS same-different task structure, and an index for learning differentiation, performance CD-CI for each participant. The learning index (i.e., difference in learning performance between CD and CI trials) was positively correlated with the neural index (i.e., differences in neural representation of same vs. different sub-structure). **F) Learning and Task structure in Lateral OFC.** There was a positive correlation between neural index of task structure representation in Lateral OFC and learning index. **G) Hippocampus and Lateral OFC represent task structure similarly.** There was a strong correlation between the hippocampus neural index and lateral OFC neural index. Error bars denote within-subjects 95 % confidence intervals.

We found that participants responded optimally on 90% (SD = 9%) of CD trials and 82% (SD = 14%) of CI trials. A direct comparison between the sub-structures showed that the proportion of optimal decisions (as defined in the learning phase section) was higher for CD than CI trials (*F*(1,21) = 4.48, *p* = .049). The near-ceiling accuracy levels for CD trials indicates that participants successfully learned to make decisions based on the context, rather than overall frequency of “like” responses (see Fig. 2B).

### Task structure representation in Hippocampus and OFC: Differentiation of task structures

Although the difference between CD and CI foods was not explicitly stated, the behavioral data described above indicate that participants successfully learned and differentiated between the two types of sub-structures during the learning and decision phase.

If the hippocampus represents the learned task structure and generalizes across the decisions that shared that structure, we would predict higher pattern similarity between pairs of CD trials (e.g., an optimal decision about the apple in Store A & an optimal decision about the carrot in Store C) and between pairs of CI foods (e.g., an optimal decision about the banana in Store A & an optimal decision about the donut in Store D), compared to the similarity between CD and CI trials (e.g., an optimal decision about the apple in Store A & an optimal decision about the donut in Store D).

To test whether the task structure was represented in a manner that generalized across the two sub-structures which were possibly differentiated during learning, we computed voxel-wise hippocampal pattern similarity (PS) (collapsed across hemispheres) for same sub-structure trial pairs (CD-CD or CI-CI) and different sub-structure trial pairs (CD-CI). Analyses of these data revealed that hippocampal PS was significantly higher for Same than Different trial pairs [significant effect of task structure; *F*(1,21) = 4.89, *p* = .038], consistent with the hypothesis that the hippocampus represents information about the task structure (see Fig. 2C and 2D).

Next, we examined the task structure representation in the OFC. Based on anatomical connectivity-based parcellations (Neubert et al., 2015), OFC can be divided into three subregions; medial, lateral, and central. Several studies provided evidence for dissociable functions of subdivisions of the OFC (Noonan et al., 2010; Rudebeck & Murray, 2011). Therefore, we separated OFC into its subdivisions and examined task structure representation with a two-way repeated measures ANOVA including task sub-structure (same, different) and OFC subdivision (lateral, medial, and central) as factors. There was an interaction between task structure and OFC subdivision (*F*(1,21) = 4.20, *p* =.037). Further analyses showed that PS values were higher for Same task structure trial pairs than for Different Task structure trial pairs in lateral (same > different, *t*(41.3) = 3.65, *p* < 0.001) OFC, whereas no significant effects were observed in medial [*t*(41.3) = 1.77, *p* = 0.084] or central [t(41.3) = 0.87, *p* = 0.39) OFC (See Fig. 2E).

### Learning performance predicts hippocampal representations

In addition to examining task structure representations in Hippocampus and OFC during the Decision Phase, we also investigated the relationship between behavioral performance during the Learning Phase on Day 1/Day 2 and neural representations in the hippocampus and OFC during the Decision Phase on Day 2. On Day 1 and the beginning of Day 2, participants implicitly learned the outcomes between each food item and each store context through trial-and-error. The underlying task structure (i.e., CD and CI foods) was not explicitly instructed, therefore participants demonstrated different learning patterns. Some individuals differentiated between CD and CI foods and showed better performance for CD foods than CI foods, while others had similar performance for both CD and CI foods. If participants learned the task structure in a way that differentiated between the two task structures (instead of just memorizing each individual event), then this differentiation might be evident in the neural representations during the Decision Phase. In other words, a greater difference in performance on CD and CI foods during learning would be predictive of a greater difference in neural representations of CD and CI foods during the Decision Phase.

To understand how individual differences in learning performance are related to neural representation of the task structure during the Decision Phase, we computed the average difference in performance between CD and CI foods from the Learning Phase on Day 1 and 2 for everyone (i.e., a learning index). We then computed the difference between neural representations of CD and CI foods in the hippocampus and OFC during the Decision Phase (i.e., a neural index). Specifically, for each participant, we calculated average of the “same sub-structure” representational values (i.e., CD/CD and CI/CI) and subtracted it from the average of “different sub-structure” representational values (i.e., CD/CI). We correlated the learning index with the neural index to examine the relationship between learning performance and neural representations of the task structure (See Fig. 2F and 2G).

Results showed that the learning index was significantly correlated with the neural index in hippocampus (*R* = 0.51, *p* = 0.016), whereas the correlation in OFC was relatively high but not statistically significant (*R* = 0.41, *p* = 0.055). The correlations showed that larger learning differences between CD and CI trials predicted stronger neural differentiation of the two sub-structures, suggesting that the learning of the task structure predicted the extent to which the structure was evident subsequently during decisions in hippocampal voxel patterns (see Fig. 2E and 2F).

### Related task structure representations in Hippocampus and OFC

We next investigated the relationship between the hippocampus and OFC in this task. If hippocampus and OFC both represent the task structure, then we reasoned that task structure representation indices should be similar for the two regions across subjects. For each participant, we calculated average of the “same-sub-structure” representational values (i.e., CD/CD and CI/CI) and subtracted it from the average of “different sub-structure” representational values (i.e., CD/CI) for both hippocampus and OFC. When we correlated these values from each participant with each other, we found a striking correlation between task structure representation in the hippocampus and lateral OFC: those individuals with stronger representations of task structure in hippocampus also had stronger representations of task structure in OFC (*R* = .71, *p* < .0001). This finding suggests that there was strong concurrence between hippocampal and OFC representations of task structure (see Fig. 2G).

### Item and Context Representations in Hippocampus and OFC

To further investigate what type of information hippocampus and OFC extract and represent from the task, we next examined whether hippocampal and OFC activity patterns also carry information about the specific contexts in which the decisions were made and/or the items the decisions were made about. There are at least two ways in which items and/or contexts can be represented. One possibility is that hippocampus generalizes across any trials that share similar properties. That is, overlap of the food-preference mappings between stores or between foods could result in similar neural patterns. For example, foods that have the same outcome across all stores (i.e., apple and muffin) might have more similar neural patterns compared to two foods that have different outcomes across all stores (i.e., apple and carrot). There is also overlap between stores that share the same food-preference mappings across all foods, which could result in similar neural patterns between stores (e.g., stores A and B would have a similar neural pattern and stores C and D would have a similar neural pattern). Another possibility, based on findings from many recent fMRI studies (see Ritvo et al., 2019 for a review), is that overlap between item or context features could instead result in dissimilar patterns in the hippocampus, leading to greater dissimilarity between foods or stores that share similar properties.

Generalization can occur at the item level (i.e., across similar foods) or at the context level (i.e., similar stores) based on their outcomes (i.e., like vs. dislike probabilities). Given that there was evidence of sub-structure representation in both the hippocampus and OFC (i.e., CD and CI foods were differentiated), we investigated item and context representations for CD and CI foods separately.

For CD foods, we performed repeated measures ANOVAs on PS values with factors for food type (similar, different) and store type (similar, different) separately for hippocampus and OFC, and we performed the same analyses for CI foods. These analyses revealed no significant effects involving food type or store type for either hippocampus (all F’s < 2.69, all p’s > 0.11) or OFC (all F’s< 2.39, all p’s> 0.13 (see Figure S1 and tables SM-3 to SM-6for the full results in the Supplemental Material).

## DISCUSSION

The goal of the present study was to investigate the role of the hippocampus and OFC in a memory-guided decision-making task. In our experiment, participants needed to learn and integrate information about items (i.e., foods), contexts (i.e., stores), and task structures (context- determined vs. context invariant probabilistic) that could help making accurate future decisions. Even though participants were never explicitly informed about the distinction between CD and CI trials, activity patterns in the hippocampus and lateral OFC carried information about this task structure. Moreover, task structure representation in the hippocampus was behaviorally relevant, in that it predicted the extent to which participants showed behavioral differences between CD and CI trials during learning. Finally, we found a striking correlation between task structure representations in hippocampus and lateral OFC. Collectively, these findings are consistent with the hypothesis that hippocampus and OFC collectively represent task structure during decision-making.

### Representation of task structure in Hippocampus

If memories of past experiences are used to guide a decision, then the critical question is: how is decision-relevant information represented in memory? In our paradigm, there are many kinds of information that could conceivably be retrieved, some of which would be relevant for predictions, and some of which would be irrelevant. The simple “episodic memory retrieval” interpretation would be that, when making each prediction, subjects would simply recall past experiences with each item, as they would in a traditional episodic memory task (Bornstein & Norman, 2017). In other words, participants could have represented outcomes for each food in each store individually, as 32 unique events. It is conceivable that participants could have used this kind of simple memory retrieval strategy and memorized each of the 32 food-store-outcome associations separately. This account would suggest that participants would form highly differentiated representations, such that there would be no generalization across food items or contexts—every item-context pairing should be associated with a unique pattern. This prediction is based on standard models of hippocampal memory functions (Davachi, 2006; Eichenbaum et al., 2007; Ranganath, 2010), but does not fully account for the present results. Instead, we see generalization across foods that were CD or CI.

Another possibility is that the hippocampus generalizes across decisions made in particular stores (contexts), which would be expected from fMRI studies of episodic memory paradigms (e.g., Dimsdale-Zucker et al., 2018; Komorowski et al., 2013; Libby et al., 2019; Ritchey et al., 2015; Schlichting & Preston, 2015; Shohamy & Wagner, 2008; Tompary & Davachi, 2017; Zeithamova & Preston, 2010). Alternatively, the hippocampus might have generalized across items based on similar outcomes (e.g., Zeithamova et al., 2018). We were surprised to find that neither of these predictions adequately accounted for the present results. We did not find evidence that the hippocampus generalized more across foods that had the same outcomes (e.g., were liked in the same stores) than foods that had different outcomes (e.g., were liked in different stores). Instead, we found that the hippocampus represented decision-relevant information from past experiences at the level of task structure. Of course, it remains possible that such information is represented in the hippocampus, as we cannot make any inferences or conclusions from null results. Nonetheless, we did find that the hippocampus represented the task relevant information that was the task structure.

Hippocampal representations during the decision phase paralleled the structure of the task—on every trial, the participant was asked to make a prediction about a particular food in a particular store. CD and CI are abstract properties that are inferred from knowledge of the task structure. In other words, one would only generalize across CD foods and differentiate them from CI foods if they understood of this aspect of the task structure. Consistent with this idea, task structure representation was strongest for participants who showed more behavioral differentiation between CD and CI trials during initial learning (Figure 2F).

Our results add in an important way to results from other studies investigating representation of decision-relevant information in the hippocampus and medial temporal lobe cortex during decision-making (Aronov et al., 2017; Bellmund et al., 2018; Park et al., 2020). For instance, Eichenbaum and colleagues used single-unit recording studies in rats to investigate hippocampal representation of deterministic, item-context-outcome associations (McKenzie et al., 2014, 2016). Their results indicated that the hippocampus represented task information in a hierarchical manner. Other work has shown that the hippocampus and/or entorhinal cortex represent information about abstract state spaces during performance of decision-making tasks that have a two-dimensional stimulus structure (Aronov et al., 2017; Garvert et al., n.d.; Theves et al., 2019). In all these studies, the tasks were designed such that one can adopt a single strategy or model to solve the task. In our experiment, the task was structured such that optimal performance could be achieved by differentiating between CD and CI trials, and, surprisingly, this factor dominated hippocampal representations during later decision making. Thus, our results suggest that, in addition to examining representations of stimuli or spatial contexts, it might be important to manipulate task structure to fully understand hippocampal contributions to prediction and decision-making.

### Representation of task structure in OFC

In addition to the hippocampus, researchers have recently proposed that the OFC might also form cognitive maps that are used for decision making. Consistent with this idea, several findings suggest that the OFC represents the space of information that is used to make decisions (Farovik et al., 2015; Schuck et al., 2016). Farovik et al. (2015), for instance, examined how decision relevant task dimensions were represented in OFC of rats as they performed a context-guided object discrimination task. Results showed that OFC neurons sharply differentiated between events associated with reward outcomes and those that were not. Within each category of rewarded and nonrewarded outcomes, hippocampal neurons generalized across encounters with the same object in the same context.

We did not find evidence to support the idea that the OFC formed generalized representations of foods based on their probability of preference (item similarity) and instead, we found that OFC carried information about whether participants were making CD or CI decisions. There are several important differences between the study of Farovik et al. (2015) and the present study. Farovik et al. (2015) provided explicit rewards for correct performance, whereas we examined activity during trials in which participants judged customer preferences, in which participants were not presented with any feedback about the accuracy of their judgments. Another key difference is that, in Farovik et al. (2015), the correct decision was entirely dependent on the context, whereas, in our study, context only was relevant for CD trials and not for CI trials. Putting these factors together, it is likely that our task-oriented participants to focus more on task structure, rather than item specific preferences per se.

Our results align well with a model recently proposed by Zhou et al. (Zhou et al., 2021), who argue that representation of value in OFC is relevant to its more general role in representations of task structure. One prominent study which provided evidence for the role of OFC in task representation was by Schuck et al. (2016). Schuck et al. (2016) designed an fMRI study in which participants were asked to judge whether a face or a house was young or old on each trial. Images of a face and a house were spatially superimposed and were presented on the same trial. Participants were asked to only judge the face or the house on each trial and whether to judge a face or a house was determined by a change of age on the previous trial. This required participants to infer the trial type, a hidden state that needed to be learned from previous trials and created 16 unique trial types. They showed that OFC represented these task states and that it is an important region for task structure representation. This study did not have explicit values assigned to faces or images, therefore, OFC involvement in this task was not directly tied to a value representation. Other studies also provided evidence for OFC representing hidden states in a task (Park et al., 2020; Vaidya et al., 2021; Wimmer & Büchel, 2019). Sometimes these hidden states could be predictive of a reward as well (e.g., Zhou et al., 2019). In these cases, it was shown that OFC represented both value and task states but could be identified separately in OFC.

Again, this doesn’t diminish a role of OFC in value representation. It’s also possible that in tasks that have a simple structure with explicitly rewarded outcomes, OFC will predominantly represent value information (Enel et al., 2020), whereas in situations where one might adopt multiple models to guide decision-making, task structure information might be more prominent.

### Related task structure representations in Hippocampus and OFC

In the past, research on decision-making has focused on OFC, whereas research on the hippocampus has focused on memory. More recently, it has become apparent that Hippocampus and OFC interact with each other during learning and decision-making (Knudsen & Wallis, 2020; Wikenheiser et al., 2017), and it has been suggested that these regions collectively represent task structure (Wikenheiser & Schoenbaum, 2016). Medial portions of OFC receive direct inputs from the hippocampus, and available evidence suggests that the hippocampus might be a source of context information for the OFC (Eichenbaum, 2017; Navawongse & Eichenbaum, 2013). Wikenheiser and Schoenbaum (2016) suggested that it is possible hippocampus and OFC encode variables in parallel but interact with each other during making of cognitive maps, a unified representation of an environment to guide future behavior. Furthermore, one recent study showed that representation of task structure in OFC was dependent on hippocampal output (Wikenheiser et al., 2017). It has been suggested that hippocampus and OFC play complementary roles in representing task structure in which they cooperate in organizing information that guides behavior (Zhou et al., 2019).

Based on the results described above, we investigated the relationships between representation of task-relevant information in the Hippocampus and OFC. Our findings showed that lateral OFC, which receives direct projections from the hippocampal formation (Barbas & Blatt, 1995), generalized across decisions that involved a similar task structure (i.e., CD-CD and CI-CI > CD-CI), whereas this effect was not apparent in central and medial OFC. Notably, this lateral subdivision on OFC is thought to be most similar in terms of both its anatomical connectivity and function to the rodent OFC (Murray et al. 2007; Rusworth et al., 2011). We found a striking correlation between task structure representation indices of hippocampus and lateral OFC. Individuals who showed stronger representations of task structure in hippocampus showed stronger representations of decision-type in OFC. This finding suggests that there was strong concurrence between hippocampal and OFC representations of task structure. Thus, our results align with accumulating evidence showing parallel development of task structure representations in the Hippocampus and OFC.

In our task, Hippocampus and OFC represented the task structure and no other task related information. However, this does not necessarily imply that the two regions capture the same information from the task. If we had introduced explicit rewards or if we tracked representational similarity during the learning phase, we might have observed differences between the two regions. Additionally, “task structure” involved the extent to which food outcomes were both context-dependent and deterministic, or context-independent and probabilistic. Future studies can separately manipulate context dependence and the type of contingency (probabilistic vs. deterministic) to address the extent to which these factors might be represented by different brain regions, including Hippocampus and OFC.

### General Conclusion

In summary, the results suggest that, after learning, hippocampus and OFC carry information about the task structure, presumably the most relevant to decisions. Additionally, we demonstrate a relationship between HC and OFC in representing the task-relevant abstracted structure and the extent to which people’s behavior used this information. Our findings add to a growing body of literature showing that goal-relevant information is extensively represented in the hippocampus and the task structure is jointly represented in hippocampus and OFC.

## MATERIALS AND METHODS

### Participants

Thirty-five healthy young adults (female=20; mean age = 22. 18 years, *SD* = 3.85 years) without any neurological or psychiatric disorders were recruited from the University of California, Davis Psychology Department subject pool. All thirty-five participants completed the behavioral learning session, however, eight participants performed below the set criteria for learning (60 % performance) and did not participate in the second session. One participant did not complete the second session due to a technical issue and another participant who achieved criterion level performance was unable to take part in the second session. The remaining 25 participants completed the second MRI scanning session which included a learning, decision, and a choice phase. Participants with head motion greater than 3 mm from origin (N=3) were excluded from the analysis, leaving 22 participants (female = 12) who completed both sessions and were included in the analyses reported here. Participants were compensated with either a check or an amazon gift card. All procedures were approved by the University of California, Davis Institutional Review Board.

### Stimuli and materials

Stimuli consisted of 8 food categories items and 4 grocery store scenes. Grocery store images were the same across all participants. Participants were presented with one food item from each of the 8 categories (apple, muffin, carrot, bread, donut, onion, banana, cookie), which was randomly selected from a pool of 16 different exemplars of that food category. All participants, therefore, saw an exemplar from each of the 8 food categories, however the exact food images were different (e.g., every participant saw a banana but not every participant saw the same banana picture). Additionally, food items were randomly assigned to either “like” or “dislike” preferences in each of the four stores. This food-preference mapping was consistent within each participant but randomized across participants (e.g., for one participant banana was liked in store A but for another participant banana could be disliked in store A).

### Procedure

The experiment consisted of two sessions that took place across two consecutive days (see Figure 1 and Figure S1). During session 1, participants completed the learning task outside of the scanner, where they learned which foods were preferred in which stores through repeated trials of cycles of decisions and feedback. The first eight blocks of the learning phase trained participants only in one store such that the block trials only showed one store and provided each food’s desirability in that store. The following four blocks trained participants in different stores. In mixed blocks (blocks 9 to 12), each store was trained, the order of stores was randomized. Participants completed two blocks of learning on the second day which was similar to the last four blocks of day 1 learning. Learning performance was based on the last four blocks of day 1 and the two blocks of learning on day 2.

For the first 8 learning blocks (1-8) on the store context (i.e., background of mini-block) was constant (same store for all four mini-blocks). In blocks 9-12, the store context changed with each mini-block. The order of the mini-block store contexts was randomized, such that stores 1-4 were presented in a random order in blocks 9-12. During the mini-block, participants completed 8 trials. During each trial, a store scene was presented on the screen followed and a picture of a food item appeared, overlaid on top of the of the store along with response options for “like” or “dislike.” Participants had 2 seconds to select a response using the j and k keys. After 2 seconds, the food item disappeared and the word “CORRECT” or “INCORRECT” was displayed along with a picture of a customer’s reaction to that food (i.e., a disgusted or pleased face). This feedback was presented on the screen for 2 seconds, followed by an inter-trial interval (ITI). The ITI during the learning phase on Day 1 (outside of the scanner) was fixed at 2s. The ITI on Day 2 (inside the scanner) was jittered with a range of 2-8s (mean = 4s). The participants made like/dislike judgments on all 8 foods within a mini-block, then the store context changed, and a new mini-block began. The time between each mini-block was jittered the same as the time between trials within the mini-block (jittered, mean = 4 s) and a fixation cross was presented on the screen. Upon the start of each mini-block, and the appearance of a new store context, just the image of the store was presented on the screen for 6 seconds, followed by the eight food trials. The presentation order of the foods within a mini-block was also randomized.

Participants returned the next day to complete session 2 and were scanned while they completed two additional runs of the learning phase (to ensure that the associations were well learned), two runs of the decision task, and four runs of a choice task. The fMRI analyses presented in this paper focused on the decision phase which included participants deciding whether customers preferred presented foods in a given store or not based on previous learning. The decision phase on Day 2 (inside of the scanner) followed the same task structure as blocks of the Day 2 learning phase, with the exception that the 2s feedback period was removed from the trial. After making their response, the food item disappeared from the screen, leaving only the store context on the screen until the start of the next trial with a new food image (jittered ITI, mean 4s).

During the decision phase, participants completed trials where a grocery store appeared on the screen followed by a food item, which was overlaid on top of the store. Using what they had learned the previous day, participants had 2 seconds to predict what the customer reactions were to that food item in that store by selecting either “like” or “dislike” using the j and k keys. Participants were not given any feedback about their answer (they did receive feedback in the learning trials). There was an inter-trial interval time between the offset of the decision of one trial to the onset of the next trial which was jittered and lasted between 2-8s (mean = 4s). Participants completed two runs of the decision phase, where each run consisted of 8 mini-blocks. Each mini-block had 8 trials (so every run had 64 trials). During the mini-blocks, the background image of the store remained on the screen for the entire mini-block and changed to a new store after each mini-block (e.g. participants visited all four stores twice in every full run, each store was a mini-block). Participants made like/dislike judgments on all 8 foods in each mini-block. The time between each mini-block was jittered (mean = 4 seconds). Each mini-block started with the presentation of the store for 6 seconds, followed by the eight foods. The order of the foods was randomized within each mini-block. The mini-blocks following each other always presented a different store.

#### “Like” probabilities

The food item and store context pairs varied in their “like” probability so that there were context-determined (CD) and context-invariant (CI) foods. Foods with “like” probability of 1 or 0 depending on the store they were presented in are considered CD, while the foods whose “like” probability are .75 or .25 across all stores are CI. There are also two stores that share the same “like” preference across all 8 foods, so these are considered as “similar contexts.” The food preference probability matrix is shown in Figure 2. The mean like preference of all foods presented within a store was .5 such that no store had more favorable preferences (probability of a row in the matrix) than another store. CI foods had the same “like” preference probability across all stores and the average probability of them was either .75 or .25 (mean probability of a CI food column). CD foods on the other hand were 100% liked in half of the stores and 0% liked in the other half of the stores. In order to predict the like preference of the CD foods, participants had to understand the relationship between the store and food preference. The overall average like probability of CD foods was .5 (mean probability of a CD food column). This probability did not provide any information about the likeability of that food across all the stores.

### Image Acquisition and Preprocessing

Scanning was performed on a Siemens Skyra 3T scanner system with a 32-channel head coil at the UC Davis Facility for Integrative Neuroscience. High-resolution T1-weighted structural images were acquired using a magnetization-prepared rapid acquisition gradient echo (MPRAGE) pulse sequence (1 mm^3^ voxels; matrix size=256 x 256; 208 slices). An additional T1-weighted image with only sagittal oriented slices was used for aligning the field of view box in the subsequent functional scans (i.e., the box was adjusted for each participant to make sure that the temporal lobes were not cut off and as much of the brain as possible was in the box). Functional images were acquired over eight runs using a whole-brain multiband gradient echo planar imaging (EPI) sequence (TR = 1220 ms; TE = 24 ms; flip angle = 67°; FOV = 192 mm; multi-band factor = 2; 38 interleaved slices; voxel size = 3 mm^3^; image matrix = 64 × 64).

Processing of the fMRI data was carried out using FEAT (FMRI Expert Analysis Tool) Version 6.00, part of FSL (FMRIB’s Software Library, www.fmrib.ox.ac.uk/fsl). The two functional scans from the decision phase underwent the following preprocessing steps: (1) Skull stripping of the anatomical images was carried out using FSL’s BET function (Smith, 2002). (2) Motion correction was carried out using FSL’s MCFLIRT function (Jenkinson et al., 2002), whereby volumes in the functional scans were aligned to the center volume of each scan with rigid body registration. (3) Grand-mean intensity normalization of the entire 4D dataset by a single multiplicative factor. (4) A high pass filter cut off was set at 100 and a high pass temporal filter was applied to remove low frequency noise from the signal (Gaussian-weighted least-squares straight line fitting, with sigma=50.0s). (5) Functional scans were coregistered to each subject’s native space anatomical scan using the boundary-based registration (BBR) (Jenkinson et al., 2002; Jenkinson & Smith, 2001) cost function with FLIRT.

### ROI definition and masks

Bilateral ROI masks for the hippocampus were manually segmented on the high-resolution anatomical using the guidelines supplied by Frankó et al. (2014). Masks for the OFC were labeled with FreeSurfer (Desikan et al., 2006; Fischl, 2004). T1-weighted anatomical scans were preprocessed using FreeSurfer which included intensity normalization, removal of non-brain tissue, transformation to Talaraich space, and segmentation of gray matter, white matter, and CSF. Surfaces were calculated for the white matter-gray matter, and gray matter-pial interface. Surface-based registration to the HCP-MMP1.0 atlas (Glasser et al., 2016) was performed and subject-specific cortical regions were calculated according to atlas boundaries. Surface-based cortical regions were converted to volumetric regions of interest and transformed into functional native space.

All ROIs were co-registered to the example functional image of the participant in FSL by using the FLIRT function and applxfm option applying the same parameters that were used to co-register their anatomical image with the functional images and pattern similarity analyses were done with the co-registered ROI masks.

### fMRI data analysis

#### Pattern similarity analysis

Multivoxel pattern similarity analyses was performed on the fMRI data from the decision task runs. The analyses were conducted on unsmoothed functional images in the native space and the EPI timeseries underwent motion correction and high-pass filtering (0.01 Hz) in FMRIB’s Software Library (FSL).

For each trial, a single beta image was estimated by single trial models, based on the approach called Least Squares Single (LSS), for event-related blood oxygenation level-dependent (BOLD) signal change, controlling for signal change due to all other trials and motion artifact, using ordinary least squares regression, resulting in 128 single-trial beta images (Mumford, Turner, Ashby, & Poldrack, 2012; Mumford et al. 2014). Parameter estimates for each trial were computed using a general linear model, with the first regressor as a stick function placed at the onset of each trial and a second regressor containing all the other trials.

Single-trial beta images from run 2 were coregistered with single-trial beta images from run 1 using FSL’s FLIRT linear registration software (6 degrees of freedom). Coregistered single-trial beta images with atypically high mean absolute z-score (based on the distribution of beta estimates for each grey matter voxel across all trials) were excluded from further analysis. Based on a mean absolute z threshold of 1.5, between 0 and 10 trials were excluded per subject with a median of 4. Beta images went through a second visual inspection to make sure all the deviant trials were excluded. This noise trial exclusion procedure is adopted from the previous pattern similarity studies in the lab (Libby et al., 2019).

The representational similarity analyses were then conducted using the RSA toolbox by Nili et al.(2014). For each region of interest (HP and OFC) and subregions of OFC (medial, lateral, central OFC), all trial patterns were correlated with each other using Pearson’s r resulting in a 128 * 128 pattern similarity matrix for each ROI. From these matrices, we extracted each pairwise correlation and made a data frame with the following variables: 1) Run, 2) Food, 3) Store, 4) Store Similarity (similar, different), 5) Food Similarity (similar, different), 6) Task structure (both CD, both CI, mix), 7) Task structure Similarity (same, different), 8) Response (both like, both dislike, mix), 9) Response Similarity (same, different), 10) Performance (both optimal, both suboptimal, mix), 11) Beta Quality (good vs. bad). The data frame for PS analyses was first filtered to exclude pairwise combinations that consisted of a trial with bad quality beta.

Our PS analyses included trial pairs that consisted of trials from the different runs (between-run analysis) excluding trial pairs from the same run. This was done to prevent any temporal autocorrelations trials within the same run could have. The noise from two different runs is unlikely to be correlated, therefore, this method prevents possible inflations in PS values (Dimsdale-Zucker & Ranganath, 2018). Additionally, the presented analyses included only the pairwise combinations in which the participant gave the “optimal” response to both trials. The exclusion was done with the filter function from dplyr package in R. The variables for filtering were: 1) Run == “different”, 2) Performance == “bothoptimal”, 3) Food == “different”. We excluded same food trials because a food could either be CD or CI and not both. If we included the pairwise combinations of same foods, this could affect task structure similarity because these combinations would only be included in the same task structure condition and not in the different one. The analyses scripts and data frames are available on OSF. After the exclusion process, the remaining pairwise combinations were then passed to repeated measures of ANOVAs conducted for each ROI.

We examined whether the similarity between trials that required same motor response was a source of pattern similarity in hippocampus and OFC. We did not find any evidence for the motor response similarity resulting in higher pattern similarity in both regions. The detailed results from these analyses are presented in the Supplemental Material Pattern Similarity Results section.

## Acknowledgments

This project was supported by a NIA 1R03AG063224-01 to C.R., ONR Grant N00014-15-1-0033 to C.R.

## Author Contributions

Conceptualization, L.A.L. and C.R.; Methodology, L.A.L, E.M., and C.R.; Investigation, L.A.L, E.M., and N.R.B; Writing – Original Draft, E.M..; Writing – Review & Editing, E.M, C.R, E.B., and, N.R.B.; Funding Acquisition, C.R.; Supervision, C.R.

## References

Aronov, D., Nevers, R., & Tank, D. W. (2017). Mapping of a non-spatial dimension by the hippocampal–entorhinal circuit. Nature, 543(7647), 719–722. https://doi.org/10.1038/nature21692

Barbas, H., & Blatt, G. J. (1995). Topographically specific hippocampal projections target functionally distinct prefrontal areas in the rhesus monkey. Hippocampus, 5(6), 511–533. https://doi.org/10.1002/hipo.450050604

Behrens, T. E. J., Muller, T. H., Whittington, J. C. R., Mark, S., Baram, A. B., Stachenfeld, K. L., & Kurth-Nelson, Z. (2018). What Is a Cognitive Map? Organizing Knowledge for Flexible Behavior. Neuron, 100(2), 490–509. https://doi.org/10.1016/j.neuron.2018.10.002

Bellmund, J. L. S., Gärdenfors, P., Moser, E. I., & Doeller, C. F. (2018). Navigating cognition: Spatial codes for human thinking. Science, 362(6415). https://doi.org/10.1126/science.aat6766

Biderman, N., Bakkour, A., & Shohamy, D. (2020). What Are Memories For? The Hippocampus Bridges Past Experience with Future Decisions. Trends in Cognitive Sciences, 24(7), 542–556. https://doi.org/10.1016/j.tics.2020.04.004

Bornstein, A. M., & Norman, K. A. (2017). Reinstated episodic context guides sampling-based decisions for reward. Nature Neuroscience, 20(7), 997–1003. https://doi.org/10.1038/nn.4573

Davachi, L. (2006). Item, context and relational episodic encoding in humans. Current Opinion in Neurobiology, 16(6), 693–700. https://doi.org/10.1016/j.conb.2006.10.012

Desikan, R. S., Ségonne, F., Fischl, B., Quinn, B. T., Dickerson, B. C., Blacker, D., Buckner, R. L., Dale, A. M., Maguire, R. P., Hyman, B. T., Albert, M. S., & Killiany, R. J. (2006). An automated labeling system for subdividing the human cerebral cortex on MRI scans into gyral based regions of interest. NeuroImage, 31(3), 968–980. https://doi.org/10.1016/j.neuroimage.2006.01.021

Dimsdale-Zucker, H., & Ranganath, C. (2018). Representational Similarity Analyses: A Practical Guide for Functional MRI Applications. In Handbook of in Vivo Neural Plasticity Techniques (In D. Manahan-Vaughan (Ed.)). Elsevier.

Dimsdale-Zucker, H., Ritchey, M., Ekstrom, A. D., Yonelinas, A. P., & Ranganath, C. (2018). CA1 and CA3 differentially support spontaneous retrieval of episodic contexts within human hippocampal subfields. Nature Communications, 9(1), 294. https://doi.org/10.1038/s41467-017-02752-1

Eichenbaum, H. (2017). Time (and space) in the hippocampus. Current Opinion in Behavioral Sciences, 17, 65–70. https://doi.org/10.1016/j.cobeha.2017.06.010

Eichenbaum, H., Yonelinas, A. P., & Ranganath, C. (2007). The medial temporal lobe and recognition memory. Annual Review of Neuroscience, 30, 123–152. https://doi.org/10.1146/annurev.neuro.30.051606.094328

Enel, P., Wallis, J. D., & Rich, E. L. (2020). Stable and dynamic representations of value in the prefrontal cortex. ELife, 9, e54313. https://doi.org/10.7554/eLife.54313

Farovik, A., Place, R. J., McKenzie, S., Porter, B., Munro, C. E., & Eichenbaum, H. (2015). Orbitofrontal Cortex Encodes Memories within Value-Based Schemas and Represents Contexts That Guide Memory Retrieval. Journal of Neuroscience, 35(21), 8333–8344. https://doi.org/10.1523/JNEUROSCI.0134-15.2015

Fischl, B. (2004). Automatically Parcellating the Human Cerebral Cortex. Cerebral Cortex, 14(1), 11–22. https://doi.org/10.1093/cercor/bhg087

Frankó, E., Insausti, A. M., Artacho-Pérula, E., Insausti, R., & Chavoix, C. (2014). Identification of the human medial temporal lobe regions on magnetic resonance images: Human Medial Temporal Lobe Landmarks. Human Brain Mapping, 35(1), 248–256. https://doi.org/10.1002/hbm.22170

Garvert, M. M., Dolan, R. J., & Behrens, T. E. (n.d.). A map of abstract relational knowledge in the human hippocampal–entorhinal cortex. ELife, 6, e17086. https://doi.org/10.7554/eLife.17086

Horner, A. J., Bisby, J. A., Bush, D., Lin, W.-J., & Burgess, N. (2015). Evidence for holistic episodic recollection via hippocampal pattern completion. Nature Communications, 6, 7462. https://doi.org/10.1038/ncomms8462

Jenkinson, M., Bannister, P., Brady, M., & Smith, S. (2002). Improved Optimization for the Robust and Accurate Linear Registration and Motion Correction of Brain Images. NeuroImage, 17(2), 825–841. https://doi.org/10.1006/nimg.2002.1132

Knudsen, E. B., & Wallis, J. D. (2020). Closed-Loop Theta Stimulation in the Orbitofrontal Cortex Prevents Reward-Based Learning. Neuron, 106(3), 537–547.e4. https://doi.org/10.1016/j.neuron.2020.02.003

Komorowski, R. W., Garcia, C. G., Wilson, A., Hattori, S., Howard, M. W., & Eichenbaum, H. (2013). Ventral hippocampal neurons are shaped by experience to represent behaviorally relevant contexts. The Journal of Neuroscience: The Official Journal of the Society for Neuroscience, 33(18), 8079–8087. https://doi.org/10.1523/JNEUROSCI.5458-12.2013

Libby, L. A., Reagh, Z. M., Bouffard, N. R., Ragland, J. D., & Ranganath, C. (2019). The Hippocampus Generalizes across Memories that Share Item and Context Information. Journal of Cognitive Neuroscience, 31(1), 24–35. https://doi.org/10.1162/jocn_a_01345

Manns, J. R., & Eichenbaum, H. (2009). A cognitive map for object memory in the hippocampus. Learning & Memory (Cold Spring Harbor, N.Y.), 16(10), 616–624. https://doi.org/10.1101/lm.1484509

McKenzie, S., Frank, A. J., Kinsky, N. R., Porter, B., Rivière, P. D., & Eichenbaum, H. (2014). Hippocampal Representation of Related and Opposing Memories Develop within Distinct, Hierarchically Organized Neural Schemas. Neuron, 83(1), 202–215. https://doi.org/10.1016/j.neuron.2014.05.019

McKenzie, S., Keene, C. S., Farovik, A., Bladon, J., Place, R., Komorowski, R., & Eichenbaum, H. (2016). Representation of memories in the cortical– hippocampal system: Results from the application of population similarity analyses. Neurobiology of Learning and Memory, 134, 178–191. https://doi.org/10.1016/j.nlm.2015.12.008

Moscovitch, M., Cabeza, R., Winocur, G., & Nadel, L. (2016). Episodic Memory and Beyond: The Hippocampus and Neocortex in Transformation. Annual Review of Psychology, 67(1), 105–134. https://doi.org/10.1146/annurev-psych-113011-143733

Mumford, J. A., Turner, B. O., Ashby, F. G., & Poldrack, R. A. (2012). Deconvolving BOLD activation in event-related designs for multivoxel pattern classification analyses. NeuroImage, 59(3), 2636–2643. https://doi.org/10.1016/j.neuroimage.2011.08.076

Murray, E. A., & Rudebeck, P. H. (2018). Specializations for reward-guided decision-making in the primate ventral prefrontal cortex. Nature Reviews. Neuroscience, 19(7), 404–417. https://doi.org/10.1038/s41583-018-0013-4

Navawongse, R., & Eichenbaum, H. (2013). Distinct Pathways for Rule-Based Retrieval and Spatial Mapping of Memory Representations in Hippocampal Neurons. Journal of Neuroscience, 33(3), 1002–1013. https://doi.org/10.1523/JNEUROSCI.3891-12.2013

Neubert, F.-X., Mars, R. B., Sallet, J., & Rushworth, M. F. S. (2015). Connectivity reveals relationship of brain areas for reward-guided learning and decision making in human and monkey frontal cortex. Proceedings of the National Academy of Sciences, 112(20), E2695–E2704.

Nili, H., Wingfield, C., Walther, A., Su, L., Marslen-Wilson, W., & Kriegeskorte, N. (2014). A Toolbox for Representational Similarity Analysis. PLoS Computational Biology, 10(4), e1003553. https://doi.org/10.1371/journal.pcbi.1003553

Noonan, M. P., Walton, M. E., Behrens, T. E. J., Sallet, J., Buckley, M. J., & Rushworth, M. F. S. (2010). Separate value comparison and learning mechanisms in macaque medial and lateral orbitofrontal cortex. Proceedings of the National Academy of Sciences, 107(47), 20547–20552. https://doi.org/10.1073/pnas.1012246107

O’Keefe, J., & Nadel, L. (1978). The hippocampus as a cognitive map. Clarendon Press; Oxford University Press.

Padoa-Schioppa, C., & Conen, K. E. (2017). Orbitofrontal Cortex: A Neural Circuit for Economic Decisions. Neuron, 96(4), 736–754. https://doi.org/10.1016/j.neuron.2017.09.031

Park, S. A., Miller, D. S., Nili, H., Ranganath, C., & Boorman, E. D. (2020). Map Making: Constructing, Combining, and Inferring on Abstract Cognitive Maps. Neuron, 107(6), 1226–1238.e8. https://doi.org/10.1016/j.neuron.2020.06.030

Ranganath, C. (2010). Binding Items and Contexts: The Cognitive Neuroscience of Episodic Memory. Current Directions in Psychological Science, 19(3), 131–137. https://doi.org/10.1177/0963721410368805

Ritchey, M., Montchal, M. E., Yonelinas, A. P., & Ranganath, C. (2015). Delay-dependent contributions of medial temporal lobe regions to episodic memory retrieval. ELife, 4, e05025. https://doi.org/10.7554/eLife.05025

Ritvo, V. J. H., Turk-Browne, N. B., & Norman, K. A. (2019). Nonmonotonic Plasticity: How Memory Retrieval Drives Learning. Trends in Cognitive Sciences, 23(9), 726–742. https://doi.org/10.1016/j.tics.2019.06.007

Rudebeck, P. H., & Murray, E. A. (2011). Dissociable Effects of Subtotal Lesions within the Macaque Orbital Prefrontal Cortex on Reward-Guided Behavior. Journal of Neuroscience, 31(29), 10569–10578. https://doi.org/10.1523/JNEUROSCI.0091-11.2011

Rushworth, M. F. S., Noonan, M. P., Boorman, E. D., Walton, M. E., & Behrens, T. E. (2011). Frontal Cortex and Reward-Guided Learning and Decision-Making. Neuron, 70(6), 1054–1069. https://doi.org/10.1016/j.neuron.2011.05.014

Schapiro, A. C., Turk-Browne, N. B., Botvinick, M. M., & Norman, K. A. (2017). Complementary learning systems within the hippocampus: A neural network modelling approach to reconciling episodic memory with statistical learning. Philosophical Transactions of the Royal Society B: Biological Sciences, 372(1711), 20160049. https://doi.org/10.1098/rstb.2016.0049

Schapiro, A. C., Turk-Browne, N. B., Norman, K. A., & Botvinick, M. M. (2016). Statistical learning of temporal community structure in the hippocampus. Hippocampus, 26(1), 3–8. https://doi.org/10.1002/hipo.22523

Schiller, D., Eichenbaum, H., Buffalo, E. A., Davachi, L., Foster, D. J., Leutgeb, S., & Ranganath, C. (2015). Memory and Space: Towards an Understanding of the Cognitive Map. Journal of Neuroscience, 35(41), 13904–13911. https://doi.org/10.1523/JNEUROSCI.2618-15.2015

Schlichting, M. L., & Preston, A. R. (2015). Memory integration: Neural mechanisms and implications for behavior. Current Opinion in Behavioral Sciences, 1, 1–8. https://doi.org/10.1016/j.cobeha.2014.07.005

Schuck, N. W., Cai, M. B., Wilson, R. C., & Niv, Y. (2016). Human Orbitofrontal Cortex Represents a Cognitive Map of State Space. Neuron, 91(6), 1402–1412. https://doi.org/10.1016/j.neuron.2016.08.019

Shohamy, D., & Wagner, A. D. (2008). Integrating Memories in the Human Brain: Hippocampal–Midbrain Encoding of Overlapping Events. Neuron, 60(2), 378–389. https://doi.org/10.1016/j.neuron.2008.09.023

Smith, S. M. (2002). Fast robust automated brain extraction. Human Brain Mapping, 17(3), 143–155. https://doi.org/10.1002/hbm.10062

Theves, S., Fernandez, G., & Doeller, C. F. (2019). The Hippocampus Encodes Distances in Multidimensional Feature Space. Current Biology, 29(7), 1226–1231.e3. https://doi.org/10.1016/j.cub.2019.02.035

Tolman, E. C. (1948). Cognitive maps in rats and men. Psychological Review, 55(4), 189–208. https://doi.org/10.1037/h0061626

Tompary, A., & Davachi, L. (2017). Consolidation Promotes the Emergence of Representational Overlap in the Hippocampus and Medial Prefrontal Cortex. Neuron, 96(1), 228–241.e5. https://doi.org/10.1016/j.neuron.2017.09.005

Vaidya, A. R., Jones, H. M., Castillo, J., & Badre, D. (2021). Neural representation of abstract task structure during generalization. ELife, 10, e63226. https://doi.org/10.7554/eLife.63226

Wallis, J. D. (2007). Orbitofrontal Cortex and Its Contribution to Decision-Making. Annual Review of Neuroscience, 30(1), 31–56. https://doi.org/10.1146/annurev.neuro.30.051606.094334

Wallis, J. D. (2019). Chapter 15—Reward. In M. D’Esposito & J. H. Grafman (Eds.), Handbook of Clinical Neurology (Vol. 163, pp. 281–294). Elsevier. https://doi.org/10.1016/B978-0-12-804281-6.00015-X

Wikenheiser, A. M., Marrero-Garcia, Y., & Schoenbaum, G. (2017). Suppression of Ventral Hippocampal Output Impairs Integrated Orbitofrontal Encoding of Task Structure. Neuron, 95(5), 1197–1207.e3. https://doi.org/10.1016/j.neuron.2017.08.003

Wikenheiser, A. M., & Schoenbaum, G. (2016). Over the river, through the woods: Cognitive maps in the hippocampus and orbitofrontal cortex. Nature Reviews Neuroscience, 17(8), 513–523. https://doi.org/10.1038/nrn.2016.56

Wilson, R. C., Takahashi, Y. K., Schoenbaum, G., & Niv, Y. (2014). Orbitofrontal Cortex as a Cognitive Map of Task Space. Neuron, 81(2), 267–279. https://doi.org/10.1016/j.neuron.2013.11.005

Wimmer, E. G., & Büchel, C. (2019). Learning of distant state predictions by the orbitofrontal cortex in humans. Nature Communications, 10(1), 2554. https://doi.org/10.1038/s41467-019-10597-z

Zeithamova, D., & Preston, A. R. (2010). Flexible memories: Differential roles for medial temporal lobe and prefrontal cortex in cross-episode binding. The Journal of Neuroscience: The Official Journal of the Society for Neuroscience, 30(44), 14676–14684. https://doi.org/10.1523/JNEUROSCI.3250-10.2010

Zhou, J., Gardner, M. P., & Schoenbaum, G. (2021). Is the core function of orbitofrontal cortex to signal values or make predictions? Current Opinion in Behavioral Sciences, 41, 1–9. https://doi.org/10.1016/j.cobeha.2021.02.011

Zhou, J., Montesinos-Cartagena, M., Wikenheiser, A. M., Gardner, M. P. H., Niv, Y., & Schoenbaum, G. (2019). Complementary Task Structure Representations in Hippocampus and Orbitofrontal Cortex during an Odor Sequence Task. Current Biology, 29(20), 3402–3409.e3. https://doi.org/10.1016/j.cub.2019.08.040

