## Supplemental Material for "Representation of Task Structure in Human Hippocampus and Orbitofrontal Cortex"

**Methods – Participant performance payment**

At the end of each run of each phase, participant performance on a scale from 0 to 1 was displayed in the upper left-hand corner of the screen (0.5 being chance performance). Participants were told that their performance on the last two runs was going to be averaged and that they would be rewarded with extra bonus money based on this average. This performance was calculated for the two different trial types separately – context-determined and context-invariant – to assure that participants were learning both types of trials above chance. Therefore, there were two different numbers displayed in the top corner of the screen. Participants were not told what the two different numbers meant and were just told it was a reflection of their performance). After the 12$^{th}$ run, participant performance was averaged from the last two runs and they were rewarded if their performance was above chance ($2 for every .1 increment over .5, e.g., .6 = $2, .7= $4, .8=$6, .9=$8, and 1.0=$10). This payment was added to their overall payment at the end of the experiment ($10/hr for participating on day 1 and $20/hr on day 2). Participants moved on to Day 2 if their average accuracy from the last two runs was 60% or above (for both CD and CI separately).

**Stimuli**

Stimuli consisted of 8 foods and four grocery store scenes. Grocery store images were the same across all participants. Participants were presented with the same 8 foods but the exact food images were different. For instance, every participant saw a banana but not every participant was shown the exact same picture of the banana. For each food item, the image was randomly selected from a pool of 16 different pictures of that food item. This random selection occurred for each of the 8 foods so that every participant had a random selection of food images. Additionally, the food-outcome mappings between 8 foods and 8 like outcomes within each store was assigned randomly for each participant.

**Day 1 (outside of the scanner):**

*Learning*

Participants were instructed to act as a market researcher and predict whether or not customers like or dislike foods within different stores. They were told that not all foods are liked or disliked all the time, and that they should pay attention to the store just in case it matters. Participants saw the same 8 foods in each of the four stores. They were told that at first they would be making guesses, but that overtime they should learn to predict customer preferences in the different stores. After the instructions, all participants completed the pre-familiarization phase, where they were shown the images of every store and food.

During the first 8 learning runs, the store scene remained the same throughout the entire run (disappeared and then reappeared in between mini-blocks), so that participants saw all 8 foods four times in a single store (i.e., the store scene in the four mini-blocks remains the same throughout each run in runs 1-8). In runs 9-12, each mini-block presented a different store scene (i.e., participants saw all 8 foods once in each of the four stores). At the end of every run, participant performance on a scale from 0 to 1 was displayed in the upper left-hand corner of the screen (0.5 being chance performance). Participants were told that their performance on the last two runs was going to be averaged and that they would be rewarded with extra bonus compensation based on this average. Participants moved on to Day 2 if their average accuracy from the last two runs was 60% or above.

**Day 2 (in the scanner):**

*Choice Phase*

Lastly, participants completed four runs (64 trials in each run) of the "choice" task. During the choice runs, participants were presented with two foods within one store and were asked to choose which food was more preferred in that store, either the food on the right or the food on the left. The choice runs consisted of three trial types: a) CD choice trials, CI choice trials, and c) Competition trials.

During CD choice trials, two CD foods with 1 and 0 like outcome were presented within one store. Participants were asked to select the more preferred food in that store. For these trials, there was a definitive correct response (e.g., the food with like outcome probability 1; Apple in store A vs Bread in store A, correct response = Apple). For CI choice trials, two CI foods with .75 and .25 like outcomes were presented together in one store. Participants were asked to select the more preferred food in that store (e.g. .75; Donut in store A vs Banana in store A, correct response = Donut).

Unlike the learning and decision phases, each trial in the choice phase randomly showed one of the four stores. The two foods could either be a) two CD foods one 100 % liked and the other 0 % liked in a given store contexts, or b) two CI foods one 75 % liked and the other 25 % liked. We expected participants to choose the food that was more liked in that store based on their task structure knowledge. On averarge, participants chose the liked foods over disliked foods, 88 % of the time for choices between two CD foods, and 81 % of the time for choices between two CI foods. They were slightly better at this for CD foods compared to CI foods, however, this difference was not significant, $F(1,21) = 3.73$, $p = .07$. This showed that that participants had learned about food preferences and used their knowledge about the task to guide their future decisions and choices.

During competition trials, one CD food with a 1 or 0 like outcome was presented in the same store with one CI food with .75 or .25 like outcome. For these trials, there was no definitive correct answer (1 vs. .75 or 0 vs. .25). Instead there were item-based responses or context-based responses, where item-based responses meant relying on the averaged value of the food across all of the stores and the context-based responses meant relying on the integration of both context and item values. For instance, if the participant was in Store A and was presented with Apple (like outcome =1) and Donut (like outcome = .75), an item-based response would be choosing Donut because the average value of Donut across all stores is higher than the average value of Apple across all stores. This type of response is a context-independent response as the decision did not take into account context determined value of the food. On the other hand, if the participant chose Apple over Donut, this would be a context-dependent response because Apple within that context always yielded a higher like probability.

**Image Acquisition and Preprocessing**

Scanning was performed on a Siemens Skyra 3T scanner system with a 32-channel head coil at the UC Davis Facility for Integrative Neuroscience. High-resolution T1-weighted structural images were acquired using a magnetization-prepared rapid acquisition gradient echo (MPRAGE) pulse sequence (1 $mm^3$ voxels; matrix size=256 x 256; 208 slices). An additional T1-weighted image with only sagittal oriented slices was used for aligning the field of view box in the subsequent functional scans (i.e., the box was adjusted for each participant to make sure that the temporal lobes were not cut off and as much of the brain as possible was in the box). Functional images were acquired over eight runs using a whole-brain multiband gradient echo planar imaging (EPI) sequence (TR = 1220 ms; TE = 24 ms; flip angle = 67°; FOV = 192 mm; multi-band factor = 2; 38 interleaved slices; voxel size = 3 $mm^3$; image matrix = 64 × 64).

**Behavioral Results**

**Response Time Analysis – Decision Phase**

In order to make sure, differences in response times to optimal decisions for different sub-structures was not the source of pattern similarity, we examined the response times to CD and CI trials. We conducted a one-way ANOVA with task structure as a factor. Our analysis showed that participants responded to both sub-structures in similar times in circa 1s (Mean CD = .97, Mean CI = .94, $F(1, 21) = 1.31$, $p = 0.265$).

**Pattern similarity Results**

**Motor Response Control Analyses**

We conducted a control analysis to determine whether pattern similarity in hippocampus could be driven by the similarity of motor responses between trials, independent of other decision variables. Therefore, the analysis was based on the data from between runs but with no further exclusion based on the optimality of the decisions. Evoked patterns of activity across voxels in hippocampus ROIs were correlated for each pair of trials given either a "Like" response or "Dislike" response. For each participant, a like response was always given with the same key. If motor response was a source of pattern similarity, Like-Like trial pairs (i.e., within trials with the same motor response) would be more similar to each other than Like-Dislike trial pairs. (i.e., between trials with different motor responses). Mean similarity estimates were entered into a response similarity (same response vs. different response) repeated measures ANOVA, controlling for sub-structures (CD vs. CI) and subregions in the case of OFC. Results indicated no significant main effect of response similarity and no significant interactions. See Tables S1 and S2 for OFC and Hippocampus results.

Table S1. 1-way ANOVA with response similarity for Hippocampus and task structure as factors.

| **Effect** | **df** | **MSE** | ***F*** | **ngsquare** | ***p*** |
|---|---|---|---|---|---|
| Response sim | 1, 21 | 0 | 1.47 | 0.001 | 0.24 |

Table S2.2-way ANOVA with response similarity and OFC subregions.

| **Effect** | **df** | **MSE** | ***F*** | **ngsquare** | ***p*** |
|---|---|---|---|---|---|
| Response sim | 1, 21 | 0 | 0.02 | <0.001 | 0.88 |
| OFC subregion | 1, 21 | 0 | 17.03 | <0.001 | < 0.001 |
| Response sim * OFC subregion | 1, 21 | 0 | 1.86 | <0.001 | 0.18 |

For CI foods, we conducted the same type of analyses explained above in hippocampus and OFC. We did not find any significant effects of involving food or store type in OFC or hippocampus for CI foods (see Fig. 3C and Fig 3D). Full results are provided in the Tables 1 to 4.

Table S3. 3-way ANOVA with Store Type (similar vs. different), Food Type (similar vs. different), OFC Part (lateral vs. medial) as factors for within CD.

| **Effect** | **df** | **MSE** | ***F*** | **ngsquare** | ***p*** |
|---|---|---|---|---|---|
| Food Type | 1, 21 | 0 | 0.68 | 0.002 | 0.421 |
| Store Type | 1.59, 33.29 | 0 | 2.37 | 0.002 | 0.14 |
| OFC part | 1, 21 | 0 | 8.83 | 0.074 | < 0.001* |
| Food Type * Store Type | 1, 21 | 0 | 0.56 | 0.002 | 0.461 |
| Food Type * OFC Part | 1.6, 33.63 | 0 | 0.23 | <0.001 | 0.743 |
| Store Type * OFC Part | 1.74, 36.56 | 0 | 0.75 | <0.001 | 0.464 |
| Food Type * OFC Part * Store Type | 1.55, 32.47 | 0 | 0.92 | <0.001 | 0.387 |

Table S4. 3-way ANOVA with Store Type (similar vs. different), Food Type (similar vs. different), OFC Part (lateral vs. medial) as factors for within CI.

| **Effect** | **df** | **MSE** | ***F*** | **ngsquare** | ***p*** |
|---|---|---|---|---|---|
| Food Type | 1, 21 | 0 | 2.33 | 0.01 | 0.142 |
| Store Type | 1, 21 | 0 | 0.18 | <0.001 | 0.68 |
| OFC part | 1.99, 41.78 | 0 | 9.29 | 0.08 | 0.001 |

| | | | | | |
|---|---|---|---|---|---|
| Food Type * Store Type | 1, 21 | 0 | 1.35 | 0.005 | 0.258 |
| Food Type * OFC Part | 1.88, 39.47 | 0 | 0.85 | 0.002 | 0.427 |
| Store Type * OFC Part | 1.88, 39.57 | 0 | 0.71 | <0.001 | 0.491 |
| Food Type * OFC Part * Store Type | 1.72, 36.15 | 0 | 1.38 | 0.001 | 0.264 |

Table S5. 3-way ANOVA with Store Type (similar vs. different), Food Type (similar vs. different) for Hippocampus as factors for within CD.

| **Effect** | **df** | **MSE** | ***F*** | **ngsquare** | ***p*** |
|---|---|---|---|---|---|
| Food Type | 1, 21 | 0 | 0.17 | 0.002 | 0.68 |
| Store Type | 1, 21 | 0 | 0.01 | <0.001 | 0.94 |
| Food Type * Store Type | 1, 21 | 0 | 1.53 | 0.004 | 0.23 |

Table S6. 3-way ANOVA with Store Type (similar vs. different), Food Type (similar vs. different) for Hippocampus as factors for within CI.

| **Effect** | **df** | **MSE** | ***F*** | **ngsquare** | ***p*** |
|---|---|---|---|---|---|
| Food Type | 1, 21 | 0 | 2.68 | 0.013 | 0.12 |
| Store Type | 1, 21 | 0 | 0.22 | 0.002 | 0.207 |
| Food Type * Store Type | 1, 21 | 0 | 0.21 | <0.001 | 0.7 |

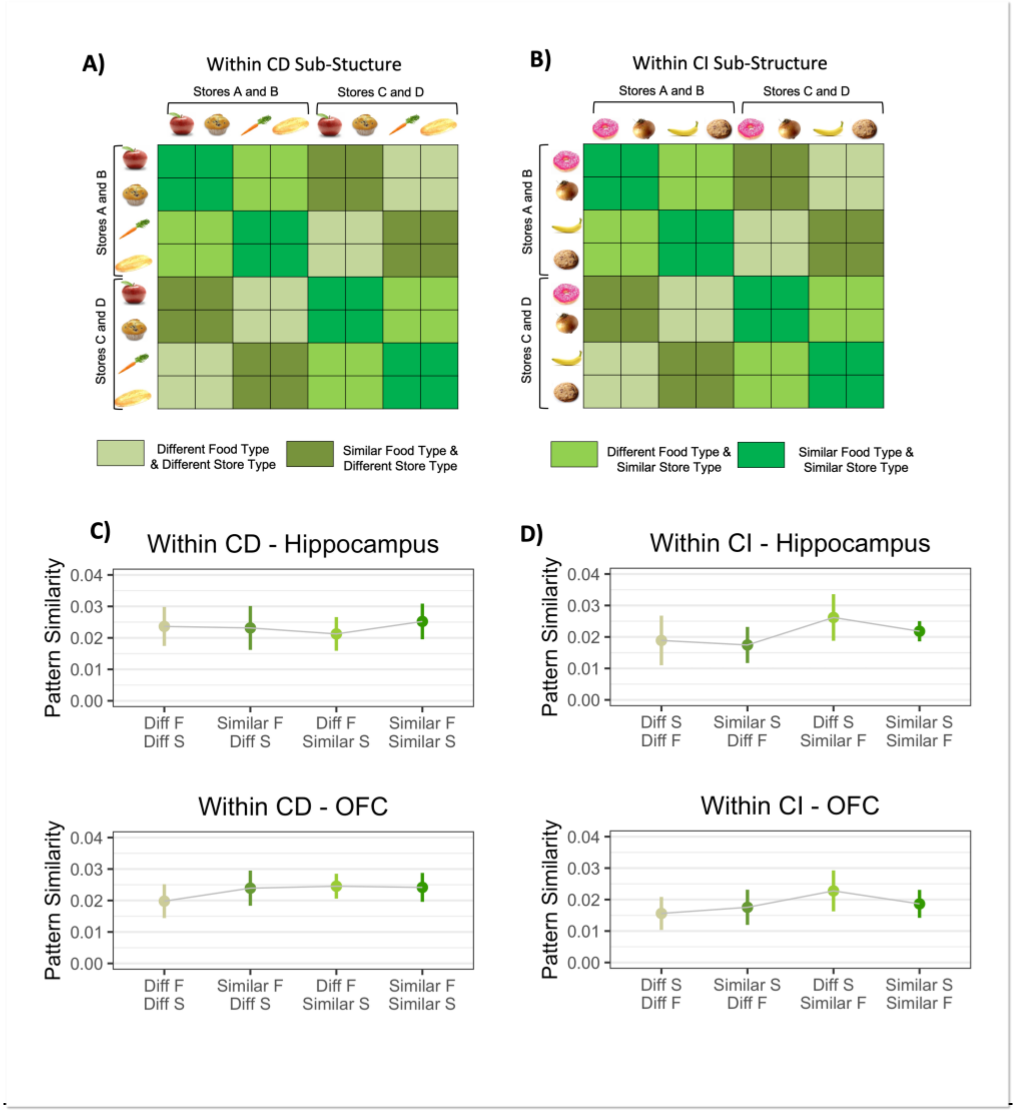


**Figure S1. A) Within CD Sub-Structure.** Item and Context similarity within CD. Color coded for different food and store types. **B) Within CI Sub-Structure.** Item and Context similarity within CI. **C) Item and Context Similarity in Hippocampus and OFC for CD Sub-Structure.** Pattern similarity values were calculated between trial pairs according to which food type and store type they were for hippocampus and OFC for the within CD. Results showed no significant variation in pattern similarity with item and context similarities. **D) Item and Context Similarity in Hippocampus and OFC for CI Sub-Structure.** Similar to CD, no significant variation in pattern similarity was observed. Diff = Different, F = Food, S = Store. Error bars denote within-subjects 95 % confidence intervals.

**Table S7**

Number of pairwise combinations for each level of task structure for OFC per participant

| | **Central OFC** | | **Lateral OFC** | | **Medial OFC** | |
|---|---|---|---|---|---|---|
| Task Structure | Same | Different | Same | Different | Same | Different |
| Participants | | | | | | |
| 1 | 5836 | 5824 | 8754 | 8736 | 5836 | 5824 |
| 2 | 4824 | 4200 | 7236 | 6300 | 4824 | 4200 |
| 3 | 5604 | 5208 | 8406 | 7812 | 5604 | 5208 |
| 4 | 5536 | 4864 | 8304 | 7296 | 5536 | 4864 |
| 5 | 6992 | 6928 | 10488 | 10392 | 6992 | 6928 |
| 6 | 5776 | 5456 | 8664 | 8184 | 5776 | 5456 |
| 7 | 4372 | 4400 | 6558 | 6600 | 4372 | 4400 |
| 8 | 4488 | 4344 | 6732 | 6516 | 4488 | 4344 |
| 9 | 5312 | 5288 | 7968 | 7932 | 5312 | 5288 |
| 10 | 6904 | 6784 | 10356 | 10176 | 6904 | 6784 |
| 11 | 7688 | 7688 | 11532 | 11532 | 7688 | 7688 |
| 12 | 6392 | 6376 | 9588 | 9564 | 6392 | 6376 |
| 13 | 5728 | 5296 | 8592 | 7944 | 5728 | 5296 |
| 14 | 7208 | 7192 | 10812 | 10788 | 7208 | 7192 |
| 15 | 6880 | 6800 | 10320 | 10200 | 6880 | 6800 |
| 16 | 6288 | 6256 | 9432 | 9384 | 6288 | 6256 |
| 17 | 5864 | 5800 | 8796 | 8700 | 5864 | 5800 |
| 18 | 6952 | 6968 | 10428 | 10452 | 6952 | 6968 |
| 19 | 6756 | 6696 | 10134 | 10044 | 6756 | 6696 |
| 20 | 7688 | 7688 | 11532 | 11532 | 7688 | 7688 |
| 21 | 5320 | 5288 | 7980 | 7932 | 5320 | 5288 |
| 22 | 7816 | 7808 | 11724 | 11712 | 7816 | 7808 |

**Table S8**

Number of pairwise combinations for each level of task structure for OFC per participant

| | **Hippocampus** | |
|---|---|---|
| Task Structure | Same | Different |
| Participants | | |
| 1 | 7752 | 7800 |
| 2 | 6000 | 5352 |
| 3 | 6264 | 5832 |
| 4 | 6954 | 6018 |
| 5 | 8148 | 8076 |
| 6 | 7992 | 7608 |
| 7 | 5550 | 5520 |
| 8 | 5730 | 5586 |

| | | |
|---|---|---|
| 9 | 5490 | 5508 |
| 10 | 7704 | 7560 |
| 11 | 9234 | 9234 |
| 12 | 8268 | 8268 |
| 13 | 6264 | 5616 |
| 14 | 8898 | 8886 |
| 15 | 9780 | 9690 |
| 16 | 7686 | 7578 |
| 17 | 6492 | 6408 |
| 18 | 9732 | 9738 |
| 19 | 8772 | 8700 |
| 20 | 9234 | 9246 |
| 21 | 5682 | 5658 |
| 22 | 10272 | 10260 |

**Table S9**

Number of pairwise combinations for each level of task structure for OFC per participant

| | Central OFC | | Lateral OFC | | Medial OFC | |
|---|---|---|---|---|---|---|
| Task Structure | CI | | CI | | CI | |
| Store Rule | Same | | Same | | Same | |
| Food Rule | Same | Different | Same | Different | Same | Different |
| Participants | | | | | | |
| 1 | 680 | 672 | 1020 | 1008 | 680 | 672 |
| 2 | 360 | 252 | 540 | 378 | 360 | 252 |
| 3 | 468 | 464 | 702 | 696 | 468 | 464 |
| 4 | 364 | 364 | 546 | 546 | 364 | 364 |
| 5 | 776 | 776 | 1164 | 1164 | 776 | 776 |
| 6 | 496 | 472 | 744 | 708 | 496 | 472 |
| 7 | 472 | 496 | 708 | 744 | 472 | 496 |
| 8 | 700 | 712 | 1050 | 1068 | 700 | 712 |
| 9 | 744 | 744 | 1116 | 1116 | 744 | 744 |
| 10 | 1024 | 1024 | 1536 | 1536 | 1024 | 1024 |
| 11 | 960 | 960 | 1440 | 1440 | 960 | 960 |
| 12 | 752 | 704 | 1128 | 1056 | 752 | 704 |
| 13 | 592 | 320 | 888 | 480 | 592 | 320 |
| 14 | 852 | 848 | 1278 | 1272 | 852 | 848 |
| 15 | 992 | 992 | 1488 | 1488 | 992 | 992 |
| 16 | 716 | 684 | 1074 | 1026 | 716 | 684 |
| 17 | 856 | 832 | 1284 | 1248 | 856 | 832 |
| 18 | 876 | 864 | 1314 | 1296 | 876 | 864 |
| 19 | 736 | 720 | 1104 | 1080 | 736 | 720 |
| 20 | 964 | 960 | 1446 | 1440 | 964 | 960 |
| 21 | 584 | 528 | 876 | 792 | 584 | 528 |

| 22 | 1024 | 1024 | 1536 | 1536 | 1024 | 1024 |
|---|---|---|---|---|---|---|

**Table S10**

Number of pairwise combinations for each level of task structure for OFC per participant

| | Central OFC | | Lateral OFC | | Medial OFC | |
|---|---|---|---|---|---|---|
| Task Structure | CI | | CI | | CI | |
| Store Rule | Different | | Different | | Different | |
| Food Rule | Same | Different | Same | Different | Same | Different |
| Participants | | | | | | |
| 1 | 680 | 672 | 1020 | 1008 | 680 | 672 |
| 2 | 276 | 336 | 414 | 504 | 276 | 336 |
| 3 | 416 | 416 | 624 | 624 | 416 | 416 |
| 4 | 356 | 356 | 534 | 534 | 356 | 356 |
| 5 | 736 | 736 | 1104 | 1104 | 736 | 736 |
| 6 | 504 | 464 | 756 | 696 | 504 | 464 |
| 7 | 440 | 464 | 660 | 696 | 440 | 464 |
| 8 | 704 | 692 | 1056 | 1038 | 704 | 692 |
| 9 | 712 | 712 | 1068 | 1068 | 712 | 712 |
| 10 | 1024 | 1024 | 1536 | 1536 | 1024 | 1024 |
| 11 | 964 | 960 | 1446 | 1440 | 964 | 960 |
| 12 | 752 | 704 | 1128 | 1056 | 752 | 704 |
| 13 | 528 | 320 | 792 | 480 | 528 | 320 |
| 14 | 832 | 832 | 1248 | 1248 | 832 | 832 |
| 15 | 992 | 992 | 1488 | 1488 | 992 | 992 |
| 16 | 680 | 728 | 1020 | 1092 | 680 | 728 |
| 17 | 844 | 832 | 1266 | 1248 | 844 | 832 |
| 18 | 876 | 864 | 1314 | 1296 | 876 | 864 |
| 19 | 736 | 720 | 1104 | 1080 | 736 | 720 |
| 20 | 960 | 960 | 1440 | 1440 | 960 | 960 |
| 21 | 576 | 512 | 864 | 768 | 576 | 512 |
| 22 | 1024 | 1024 | 1536 | 1536 | 1024 | 1024 |

**Table S11**

Number of pairwise combinations for each level of task structure for OFC per participant

| | Central OFC | | Lateral OFC | | Medial OFC | |
|---|---|---|---|---|---|---|
| Task Structure | CD | | CD | | CD | |
| Store Rule | Same | | Same | | Same | |
| Food Rule | Same | Different | Same | Different | Same | Different |
| Participants | | | | | | |
| 1 | 784 | 780 | 1176 | 1170 | 784 | 780 |

| | | | | | | |
|---|---|---|---|---|---|---|
| 2 | 912 | 896 | 1368 | 1344 | 912 | 896 |
| 3 | 960 | 960 | 1440 | 1440 | 960 | 960 |
| 4 | 1024 | 1024 | 1536 | 1536 | 1024 | 1024 |
| 5 | 992 | 992 | 1488 | 1488 | 992 | 992 |
| 6 | 960 | 960 | 1440 | 1440 | 960 | 960 |
| 7 | 620 | 624 | 930 | 936 | 620 | 624 |
| 8 | 420 | 420 | 630 | 630 | 420 | 420 |
| 9 | 652 | 660 | 978 | 990 | 652 | 660 |
| 10 | 708 | 716 | 1062 | 1074 | 708 | 716 |
| 11 | 960 | 964 | 1440 | 1446 | 960 | 964 |
| 12 | 872 | 868 | 1308 | 1302 | 872 | 868 |
| 13 | 992 | 992 | 1488 | 1488 | 992 | 992 |
| 14 | 960 | 964 | 1440 | 1446 | 960 | 964 |
| 15 | 736 | 728 | 1104 | 1092 | 736 | 728 |
| 16 | 868 | 872 | 1302 | 1308 | 868 | 872 |
| 17 | 680 | 576 | 1020 | 864 | 680 | 576 |
| 18 | 872 | 868 | 1308 | 1302 | 872 | 868 |
| 19 | 960 | 960 | 1440 | 1440 | 960 | 960 |
| 20 | 960 | 960 | 1440 | 1440 | 960 | 960 |
| 21 | 808 | 776 | 1212 | 1164 | 808 | 776 |
| 22 | 932 | 932 | 1398 | 1398 | 932 | 932 |

**Table S12**

Number of pairwise combinations for each level of task structure for OFC per participant

| | **Central OFC** | | **Lateral OFC** | | **Medial OFC** | |
|---|---|---|---|---|---|---|
| Task Structure | CD | | CD | | CD | |
| Store Rule | Different | | Different | | Different | |
| Food Rule | Same | Different | Same | Different | Same | Different |
| Participants | | | | | | |
| 1 | 784 | 784 | 1176 | 1176 | 784 | 784 |
| 2 | 896 | 896 | 1344 | 1344 | 896 | 896 |
| 3 | 960 | 960 | 1440 | 1440 | 960 | 960 |
| 4 | 1024 | 1024 | 1536 | 1536 | 1024 | 1024 |
| 5 | 992 | 992 | 1488 | 1488 | 992 | 992 |
| 6 | 960 | 960 | 1440 | 1440 | 960 | 960 |
| 7 | 628 | 628 | 942 | 942 | 628 | 628 |
| 8 | 420 | 420 | 630 | 630 | 420 | 420 |
| 9 | 544 | 544 | 816 | 816 | 544 | 544 |
| 10 | 688 | 696 | 1032 | 1044 | 688 | 696 |
| 11 | 960 | 960 | 1440 | 1440 | 960 | 960 |
| 12 | 872 | 868 | 1308 | 1302 | 872 | 868 |

| | | | | | | |
|---|---|---|---|---|---|---|
| 13 | 992 | 992 | 1488 | 1488 | 992 | 992 |
| 14 | 960 | 960 | 1440 | 1440 | 960 | 960 |
| 15 | 720 | 728 | 1080 | 1092 | 720 | 728 |
| 16 | 868 | 872 | 1302 | 1308 | 868 | 872 |
| 17 | 576 | 668 | 864 | 1002 | 576 | 668 |
| 18 | 864 | 868 | 1296 | 1302 | 864 | 868 |
| 19 | 964 | 960 | 1446 | 1440 | 964 | 960 |
| 20 | 960 | 964 | 1440 | 1446 | 960 | 964 |
| 21 | 768 | 768 | 1152 | 1152 | 768 | 768 |
| 22 | 928 | 928 | 1392 | 1392 | 928 | 928 |

**Table S13**

Number of pairwise combinations for each level of task structure for Hippocampus per participant

| | **Hippocampus** | | **Hippocampus** | |
|---|---|---|---|---|
| Task Structure | CD | | CI | |
| Store Rule | Same | | Same | |
| Food Rule | Same | Different | Same | Different |
| Participants | | | | |
| 1 | 1008 | 990 | 942 | 936 |
| 2 | 1098 | 1086 | 480 | 318 |
| 3 | 1086 | 1086 | 516 | 468 |
| 4 | 1302 | 1308 | 432 | 444 |
| 5 | 1182 | 1182 | 882 | 888 |
| 6 | 1296 | 1314 | 708 | 678 |
| 7 | 780 | 780 | 600 | 636 |
| 8 | 546 | 540 | 900 | 900 |
| 9 | 684 | 684 | 672 | 690 |
| 10 | 756 | 762 | 1176 | 1176 |
| 11 | 1140 | 1140 | 1170 | 1182 |
| 12 | 1104 | 1080 | 1008 | 948 |
| 13 | 1140 | 1134 | 570 | 294 |
| 14 | 1164 | 1182 | 1068 | 1050 |
| 15 | 1056 | 1056 | 1392 | 1392 |
| 16 | 1134 | 1128 | 786 | 768 |
| 17 | 708 | 612 | 984 | 996 |
| 18 | 1218 | 1212 | 1218 | 1200 |
| 19 | 1254 | 1260 | 936 | 918 |
| 20 | 1140 | 1158 | 1182 | 1176 |
| 21 | 786 | 786 | 696 | 564 |
| 22 | 1224 | 1230 | 1344 | 1356 |

**Table S14**
Number of pairwise combinations for each level of task structure for Hippocampus per participant

| | **Hippocampus** | | **Hippocampus** | |
|---|---|---|---|---|
| Task Structure | CD | | CI | |
| Store Rule | Different | | Different | |
| Food Rule | Same | Different | Same | Different |
| Participants | | | | |
| 1 | 996 | 1008 | 942 | 930 |
| 2 | 1098 | 1086 | 372 | 462 |
| 3 | 1074 | 1074 | 480 | 480 |
| 4 | 1302 | 1308 | 426 | 432 |
| 5 | 1170 | 1164 | 834 | 846 |
| 6 | 1314 | 1296 | 714 | 672 |
| 7 | 810 | 804 | 552 | 588 |
| 8 | 534 | 546 | 900 | 864 |
| 9 | 636 | 636 | 744 | 744 |
| 10 | 756 | 726 | 1176 | 1176 |
| 11 | 1128 | 1128 | 1182 | 1164 |
| 12 | 1104 | 1080 | 1002 | 942 |
| 13 | 1128 | 1134 | 534 | 330 |
| 14 | 1176 | 1176 | 1050 | 1032 |
| 15 | 1056 | 1044 | 1392 | 1392 |
| 16 | 1134 | 1140 | 780 | 816 |
| 17 | 660 | 612 | 954 | 966 |
| 18 | 1212 | 1218 | 1218 | 1236 |
| 19 | 1260 | 1272 | 936 | 936 |
| 20 | 1128 | 1110 | 1176 | 1164 |
| 21 | 774 | 828 | 684 | 564 |
| 22 | 1206 | 1212 | 1344 | 1356 |
